# Target the Heart: a new axis of Alzheimer’s disease prevention

**DOI:** 10.1101/2025.01.27.634057

**Authors:** Lawrence I. Heller, Allison S. Lowe, Thaís Del Rosario Hernández, Sayali V. Gore, Mallika Chatterjee, Robbert Creton

## Abstract

Cyclosporine A and other calcineurin inhibitors have been identified as prospective treatments for preventing Alzheimer’s disease. Utilizing a neural network model, Z-LaP Tracker, we previously found that calcineurin inhibitors elicit a unique behavioral profile in zebrafish larvae characterized by increased activity, acoustic hyperexcitability, and reduced visually guided behaviors. Screening a large library of FDA-approved drugs using Z-LaP Tracker revealed a cluster of 65 drugs demonstrating a cyclosporine A-like behavioral profile. 14 of these drugs were heart medications, including angiotensin receptor blockers, beta-blockers, alpha-adrenergic receptor antagonists, and a statin. This suggests some heart medications may be effective in preventing or ameliorating Alzheimer’s disease pathology. Other studies have shown that many of these 14 drugs directly or indirectly inhibit the calcineurin-NFAT pathway, alike cyclosporine A. Dual administration of the heart medications with cyclosporine A in Z-LaP Tracker revealed synergistic effects: lower doses of each heart medication could be delivered in conjunction with a lower dose of cyclosporine A to evoke a similar or larger behavioral effect than higher doses of each drug independently. This indicates that co-administering a low dose of cyclosporine A with select cardiac drugs could be a potentially effective treatment strategy for Alzheimer’s disease *and* cardiovascular dysfunction, while mitigating side effects associated with higher doses of cyclosporine A. Given that heart disease precedes Alzheimer’s disease in many patients, physicians may be able to create a treatment regimen that simultaneously addresses both conditions. Our results suggest that cyclosporine A combined with *simvastatin, irbesartan, cilostazol, doxazosin,* or *nebivolol* are the most promising candidates for future exploration.

## INTRODUCTION

Dementia is one of the greatest global medical care concerns, with over 50 million people worldwide affected today and this number expected to climb to 140 million people by 2050 {Fiest, 2016 #173}.^i^ Alzheimer’s disease (AD) is the most common form of dementia, causing 60-70% of cases {Hebert, 2013 #2}.^ii^ Treatment options and management for AD are limited. Because many AD therapies are only initiated once the patient is symptomatic, it is often too late to have an appreciable effect on AD’s most devastating symptoms. It is hypothesized that earlier treatment of relevant signaling pathways involved in neuroprotection is a better approach to preventing AD. However, the field lacks early, reliable predictors of AD.

Increases in neuronal calcium, caused by oxidative stress and accumulating Aß oligomers, over-activates calcineurin, which may be a key driver of future neural dysfunction. It is hypothesized that calcineurin mediates the neurotoxic and cognitive effects of Aß oligomers and excessive levels of activated calcineurin disrupt synaptic architecture and impairs memory {Dineley, 2010 #174}.^iii^ To this extent, elevated calcineurin levels are observed in AD patients {Norris, 2018 #3}.^iv^ Calcineurin inhibitors such as Cyclosporine A (CsA), an immunosuppressant that is FDA-approved for organ transplant rejection, have been identified as potential preventive therapeutics for AD and dementia {Reese, 2011 #4} ^v^. CsA significantly lowers incidences of dementia and AD versus the general population {Taglialatela, 2015 #5}.^vi^ While CsA may be a robust suppressor of neurodegeneration {Taglialatela, 2015 #5}, CsA and other calcineurin inhibitors have substantial side effects and immune-weakening properties. Consequently, our lab hypothesized that combining a low dose of CsA with a “CsA-similar” heart drug could mitigate side effects while bolstering the neuroprotective properties.

Using Z-LaP Tracker, a deep neural network model based on DeepLabCut {Gore, 2023 #6}, our lab has previously quantified a unique behavioral response from zebrafish larvae when exposed to CsA or other calcineurin inhibitors{Del Rosario Hernandez, 2024 #7}. ^vii^ Upon comparison with two small-molecule libraries, *Tocriscreen FDA-approved Drugs Library and Cayman Chemical FDA-Approved Drugs Screening Library*, we identified 14 FDA-approved heart therapeutics that prompted CsA-like behavioral profiles: *irbesartan, losartan, eprosartan, telmisartan, doxazosin, prazosin, nebivolol, carvedilol, simvastatin, droperidol, trifluoperazine, mirtazapine, calcifediol, and cilostazol*. These drugs directly (or two indirectly) target four receptors or pathways central to cardiac function: angiotensin receptors, alpha-1 adrenergic receptors, beta-1 adrenergic receptors, and the mevalonate pathway. A literature review revealed that all these therapies directly or indirectly inhibit the calcineurin-NFAT pathway, providing a functional link between the drug set and CsA/calcineurin inhibitors.

While cardiovascular disease (CVD) and AD share many pathologies, there have not been significant strides in identifying cardiovascular drugs or cardiovascular drug combinations that are effective in treating AD {Leszek, 2021 #8}.^viii^ In the current study, we sought to explore how dual administration of heart drugs and CsA at reduced doses would impact behavior in Z-LaP Tracker. AD and CVD are aging-related diseases. Because the aging population is likely already on a CVD medication, identifying CVD drugs that mitigate AD pathogenesis could be an effective and cost-efficient strategy for neuroprotection and cardioprotection. Furthermore, exploring already-approved heart therapies provides a shorter and safer timeline to commercialization while still preserving opportunities for patentability.

The results of the current study suggest that certain heart drugs combined with CsA may be effective in halting early stages of AD pathology. This facilitates a new treatment strategy wherein the therapy is capable of simultaneously addressing both AD and CVD. Furthermore, combining low doses of these heart drugs with low dose CsA may have synergistic efficacy effects while mitigating side effects. We hypothesize that these heart drugs and CsA may both inhibit the calcineurin-NFAT signaling pathway. The outcome of our study suggests that combining FDA-approved drugs may be a feasible alternative to developing new compounds, rapidly accelerating development time and reducing associated costs. Additionally, heart medications that have additional neuroprotective properties may be selectively prescribed to those with CVD and at risk of developing AD.

## RESULTS

### BEHAVIORAL PARAMETERS

Over 1,800 zebrafish larvae at 5 days post-fertilization were exposed to 10 µM of various heart medications, either individually or in a combination of 5 µM of each drug along with 5 µM of CsA (5 µM represents a *half-dose*). Dimethyl sulfoxide (DMSO) was used as a solvent in the stock solutions and larvae were treated with 1 µl/ml DMSO as a vehicle control. After 3 hours of exposure, larvae were shown a 3-hour PowerPoint Presentation containing visual and auditory stimuli. Throughout the presentation, the larval positions were estimated using Z-LaP Tracker, a deep neural network built upon DeepLabCut, as described in our previous publications {Gore, 2023 #6} {Del Rosario Hernandez, 2024 #7}.^ix^ ^x^ Data from the larvae positions were processed to calculate 25 behaviors for each larvae (Figure 1B). The measured behaviors include spontaneous activity levels, optomotor responses in the presence of various moving lines, and responses to acoustic stimuli with 20-second or 1-second intervals.

**FIGURE 1:**
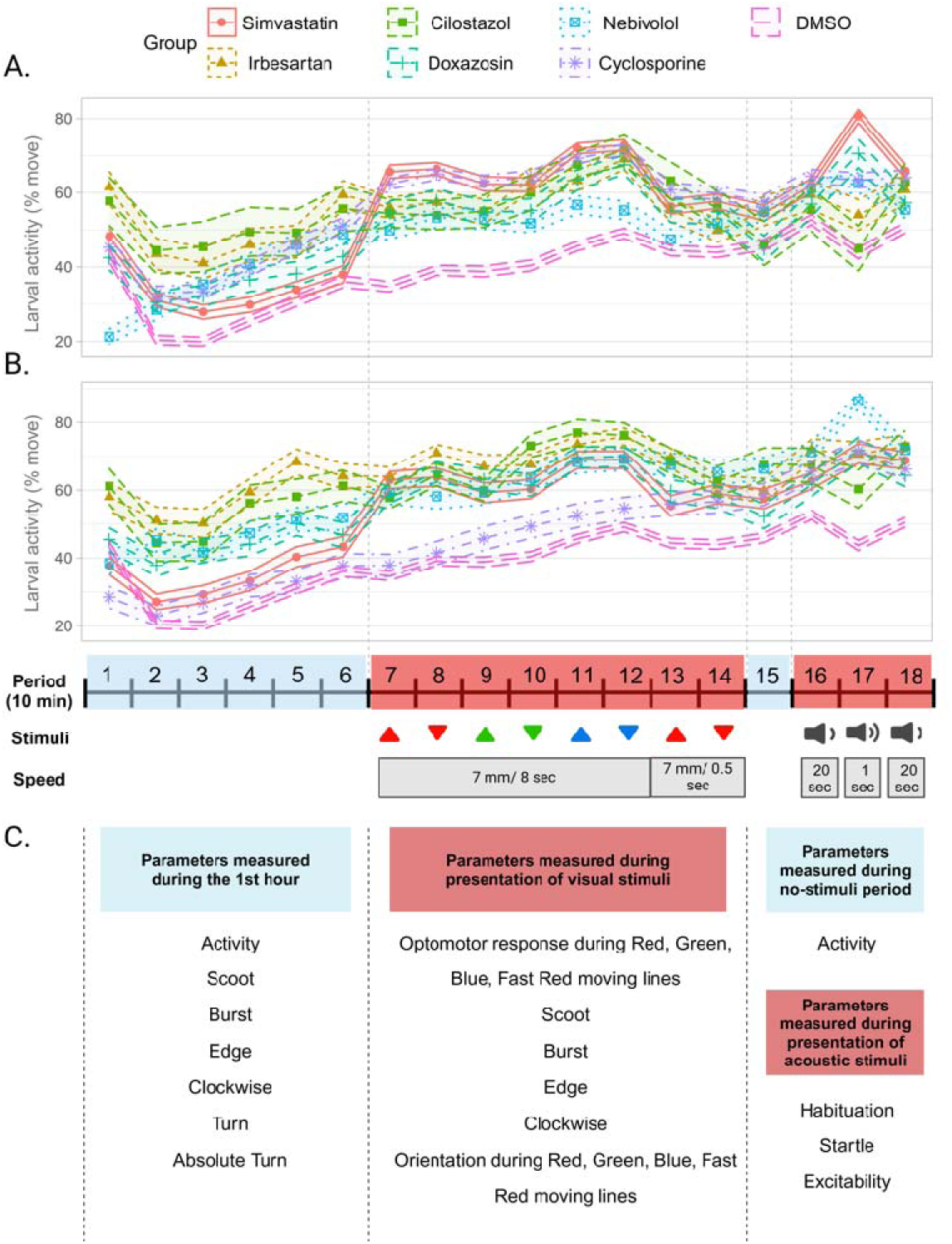
Overview of Z-LaP Tracker Presentation and Behavioral Parameters. Zebrafish larvae are shown a 3-hour presentation which includes a variety of visual and acoustic stimuli. Activity profiles of 5dpf zebrafish larvae exposed to A) 10µM and B) 5µM concentrations of the top 5 candidate drugs alone and in combination with CsA, respectively. C) Larval location is consistently measured, allowing for the computation of 25 different behavioral parameters associated with the presentation.

All 30 treatments (14 heart drugs independently, 14 heart-CsA combinations, 5 µM CsA, and 10 µM CsA) demonstrated statistically significant changes in at least one behavior compared to DMSO controls. CsA administration classically leads to an increase in baseline and resting activity, decreases in habituation, increases in excitability, decreases in optomotor responses to red moving lines, and increases in scoot and burst behaviors. Across the lower dose, dual-treatment CsA arms, CsA largely preserves this behavioral profile: increases in activity, decreases in habituation, and increases in scoot and burst behavior. Furthermore, some 5 µM combinations of CsA lead to a greater number of significant behavior changes than 5 µM CsA or the 10 µM dose of the combined drug alone, and other combinations lead to behaviors with greater changes in magnitude than 5 µM CsA or the 10 µM dose. The treatments with the greatest number of significantly changed behaviors were nebivolol, CsA 10 µM, 5 µM CsA + 5 µM nebivolol, 5 µM CsA + 5 µM simvastatin, and 5 µM CsA + 5 µM droperidol.

The 5 µM combinations resulted in differentiated behavioral profiles compared to the drugs administered individually (Figure 2A). The combination treatments were evaluated as follows: we counted the number of significantly changed behaviors using a combination treatment (e.g. 12 behaviors in the 5 µM nebivolol + 5 µM cyclosporine group) and a single treatment (e.g. 16 behaviors in the 10 µM nebivolol group) and subtracted these two values (12-16 = −4). A negative value indicates that the combination treatment is less effective than the target drug alone. A positive value indicated that the combination treatment is more effective than the target drug alone. Five 5 µM combinations had fewer statistically significant behaviors than the drug of interest alone, including nebivolol, losartan, carvedilol, telmisartan, and trifluoperazine. Seven 5 µM combinations had more statistically significant behaviors: doxazosin, irbesartan, prazosin, cilostazol, droperidol, mirtazapine, and simvastatin. Two drugs showed no change in the number of behaviors affected. The cilostazol combination led to the greatest increase in number of significantly changed behaviors.

**FIGURE 2:**
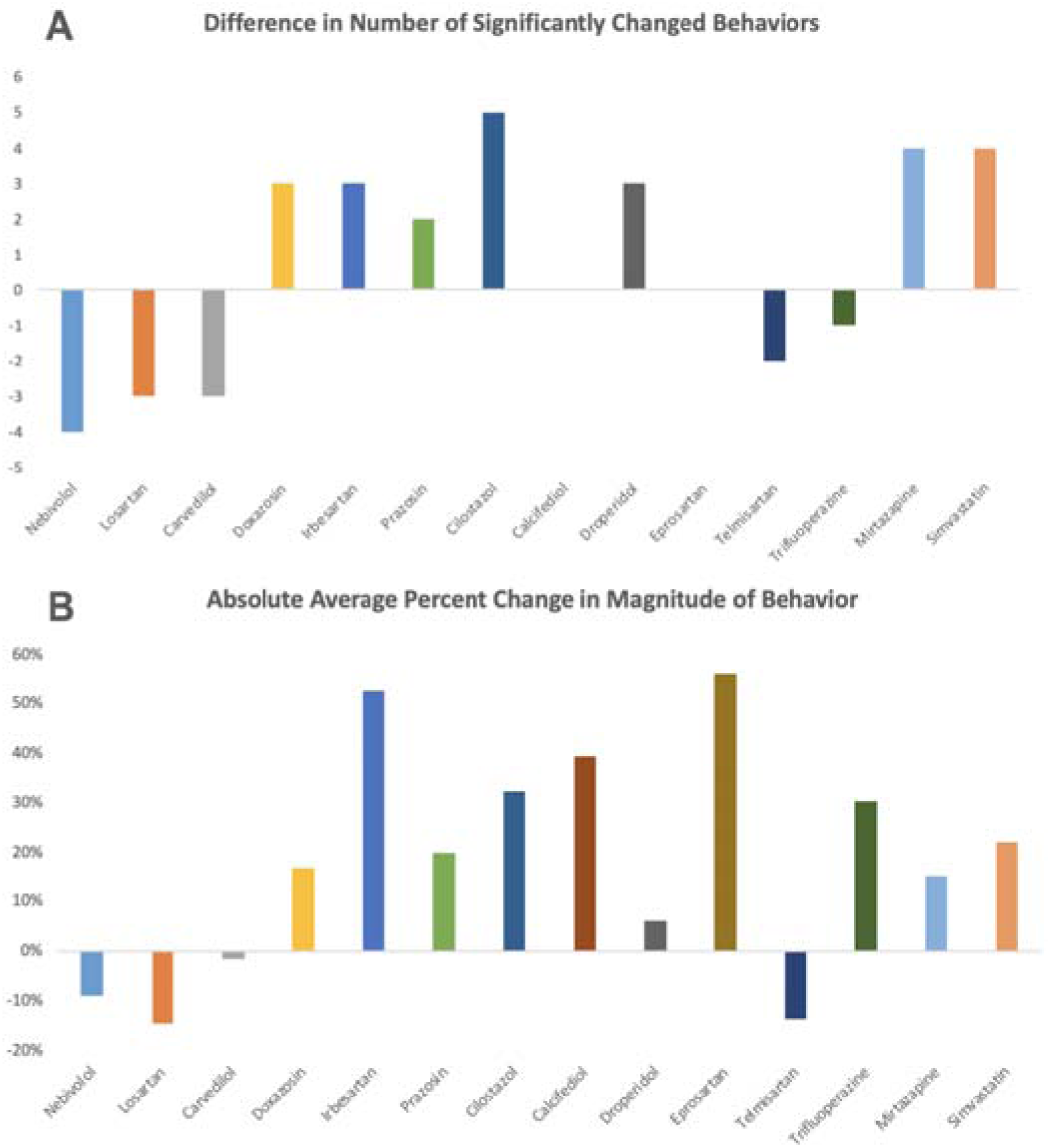
Significance and Magnitude of Behavioral Changes: The aggregated changes in the behavioral profile of each drug treatment versus its 5 µM combination with 5 µM CsA. **A)** The increase/decrease in the number of statistically significant behaviors of the 5 µM combination of each drug compared to the target drug alone. A negative number indicates that the combination treatment (e.g. 5 µM cyclosporine + 5 µM nebivolol) is less effective than the target drug alone (e.g. 10 µM nebivolol). A positive number indicates that the combination treatment (e.g. 5 µM cyclosporine + 5 µM irbesartan) is more effective that the target drug alone (e.g. 10 µM irbesartan). **B)** The percent increase/decrease in the average magnitude of behavior across all 25 observed behaviors. Again, a negative number indicates that the combination treatment is less effective than the target drug alone, and a positive number indicates that the combination treatment is more effective than the target drug alone.

Nearly all tested combinations synergized to increase the magnitude of behavior (Figure 2B). The magnitude of the effect was calculated as follows: we took the sum of the absolute average values across all behaviors of each single treatment (e.g. across all 25 behaviors, nebivolol lead to an absolute average 10.71-point change in behavior) and combination treatment (9.76-point change in behavior), and then calculated the percent change from the single treatment to combination treatment (−0.95% / 10.71% = - 8.9%). The two combinations with the largest effect were 5 µM irbesartan + 5 µM CsA, and 5 µM eprosartan + 5 µM CsA, at 53% and 56% respectively. Most combinations that led to fewer statistically significant behavioral changes also decreased the average magnitude of behavior.

### CLUSTER ANALYSIS

We employed Cluster 3.0 and Java TreeView to group the observed behavioral profiles. Hierarchical clustering was computed using Euclidean distance and complete-linkage method (Figure 3). Red and green values indicate an increase and decrease in a behavioral parameter compared to DMSO. Two relevant subclusters were identified: Subcluster A (light blue) and Subcluster B (orange). Subcluster A has a correlation value of 0.84 and includes 5 µM CsA, 10 µM CsA, and CsA combinations with nebivolol, doxazosin, and simvastatin. Within Subcluster A, 10 µM CsA, 10 µM simvastatin, and 5 µM CsA + 5 µM simvastatin yielded behavioral profiles with high correlation at 0.93. Subcluster B (correlation value of 0.85) contains combinations of CsA with irbesartan, cilostazol, calcifediol, and eprosartan; however, it does not cluster directly to CsA. While Subclusters A and B are both CsA-like (increased excitability, decreased habituation, increased scoot and burst behaviors), Subcluster A contains agents that decreased orientation to optomotor responses and had no large effect on absolute turn angle, and drugs Subcluster B generally increased optomotor response, and increased absolute turn angle with high magnitude and significance. Outside of Subclusters A and B is the DMSO Subcluster, containing prazosin, eprosartan, losartan, telmisartan, trifluoperazine, and mirtazapine. These drugs displayed low significance with CsA, with some causing decreases in activity, and others eliciting decreases in excitability.

**FIGURE 3:**
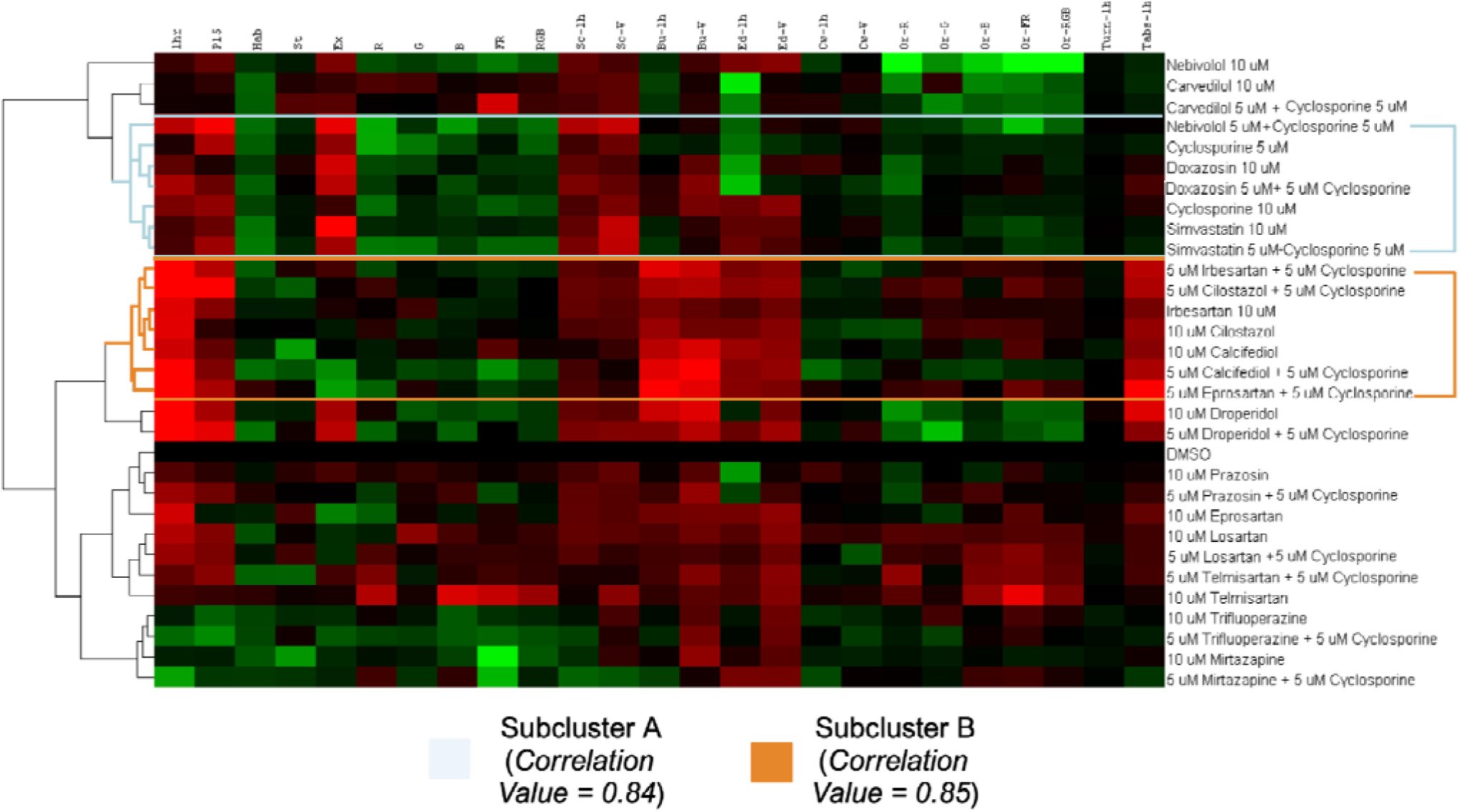
Hierarchical Clustering Analysis: Cluster TreeView of 25 behavioral parameters across 16 compounds, 14 combinations, and a DMSO control. Subcluster A demonstrates a CsA-like behavioral profile. A red box indicates an increase in a particular behavioral parameter (on x-axis) and a green box indicates a decrease. Higher color intensity represents a greater magnitude of change.

Figure 4 reveals the drugs and drug combinations that elicit the most similar changes in behavior to CsA. 5 µM CsA + 5 µM nebivolol, 5 µM CsA + 5 µM irbesartan, 5 µM CsA + 5 µM doxazosin, and simvastatin and 5 µM CsA + 5 µM simvastatin demonstrated the highest levels of positive correlation to CsA. Furthermore, the matrix displays which drugs displayed the most correlation to its 5 µM combination with 5 µM CsA. Interestingly, 5 µM simvastatin, droperidol, carvedilol, telmisartan, and trifluoperazine combined with CsA to elicit behavior most similar to higher doses of each drug alone, suggesting combinatory synergies.

**FIGURE 4:**
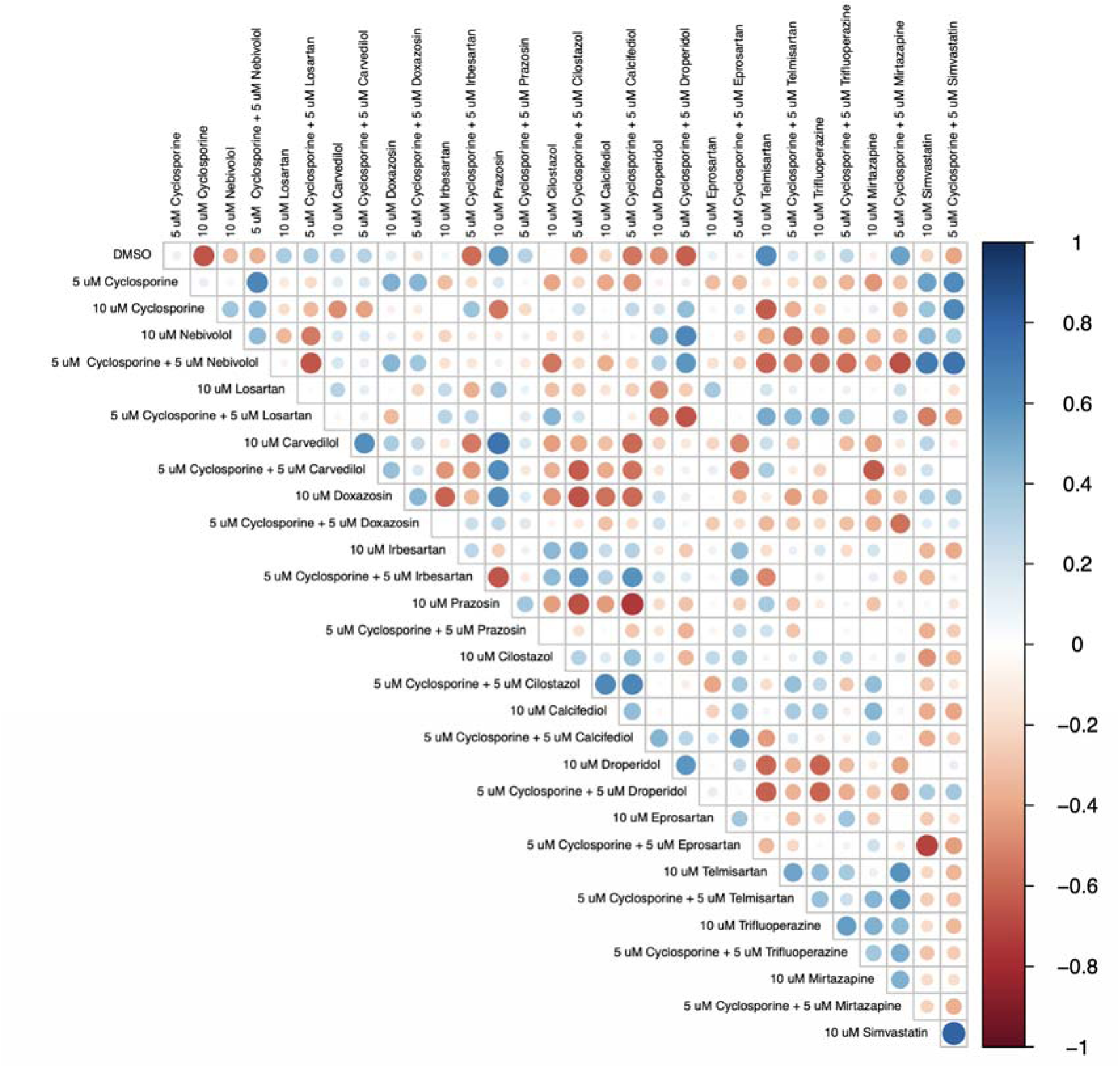
Correlation Matrix: Correlation analysis for evaluating the relationship between all 30 treatments. The strength of the relationship is denoted by a circle size and color intensity. A blue circle indicates a positive correlation and red represents a negative relationship.

### INGENUITY PATHWAY ANALYSIS

Using Ingenuity Pathway Analysis (IPA), we explored direct and indirect connections between the CsA-like heart drugs and AD. We created four separate pathways for each class of heart medications explored: alpha adrenergic receptor antagonists (doxazosin, prazosin, droperidol, trifluoperazine, and mirtazapine) (Figure 5A), beta blockers (nebivolol and carvedilol) (Figure 5B), angiotensin receptor blockers (irbesartan, losartan, eprosartan, telmisartan, calcifediol, and cilostazol) (Figure 5C), and simvastatin, an HMG-CoA reductase inhibitor (Figure 5D). Connections were overlaid using the Molecule Activity Prediction (MAP) feature, upregulating the drugs of interest. Using IPA’s *Organic* feature, the network was organized where proximity and clustering denote the strength of the relationships. These figures show that simvastatin upregulation yields high predicted inhibition of AD, beta blockers and alpha-adrenergic antagonists cause moderate predicted inhibition of AD, and that angiotensin receptor blockers have no predicted effect on AD. The beta blockers have 46 “Paths” or connections to AD, the angiotensin receptor blockers have 172 paths to AD, alpha adrenergic antagonists have 90 paths, and simvastatin has 112 paths connecting to AD.

**FIGURE 5:**
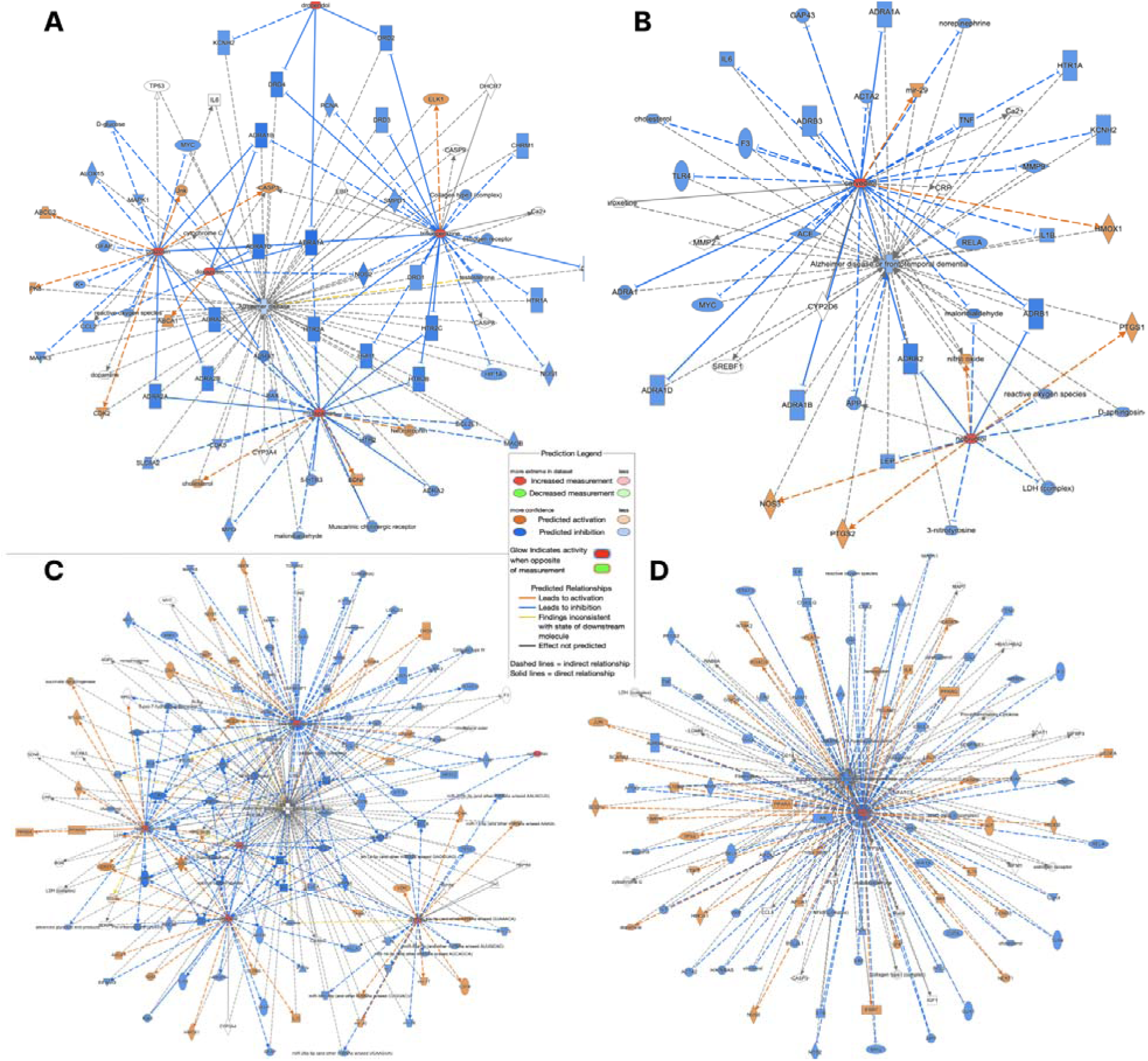
Ingenuity Pathway Analysis: The subset of 14 heart medications were divided into four categories: alpha adrenergic antagonists (A), beta blockers (B), angiotensin receptor blockers (C), and a statin (D). Each category of molecules was then connected to AD using IPA’s Path Explorer. The orange targets and lines indicate predicted activation, while blue targets and lines represent predicted inhibition. The color intensity represented the confidence of the signal. The drugs of interest are colored in red, highlighting their activation. Dashed lines indicate indirect relationships, and solid lines indicate direct relationships.

**FIGURE 6:**
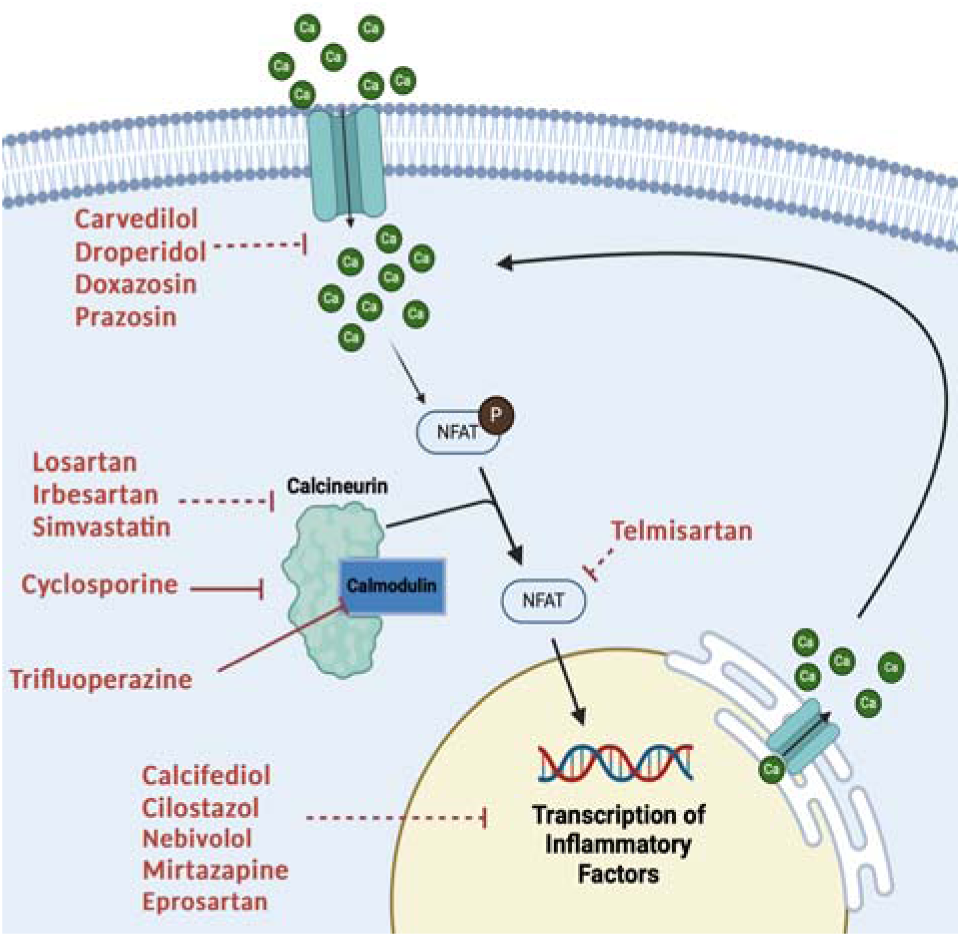
Calcineurin-NFAT Pathway: Increased intracellular calcium stimulates the calcium ion-binding protein calmodulin, which binds and activates calcineurin protein phosphatase. Calcineurin dephosphorylates proteins such as NFAT, which promotes transcription of inflammatory factors. While CsA (Cyclosporine) inhibits calcineurin, the other heart medications explored in this study also directly or indirectly inhibit fundamental molecules in this pathway. Indirect effects are indicated with dotted lines. Image created in BioRender.

**FIGURE 7:**
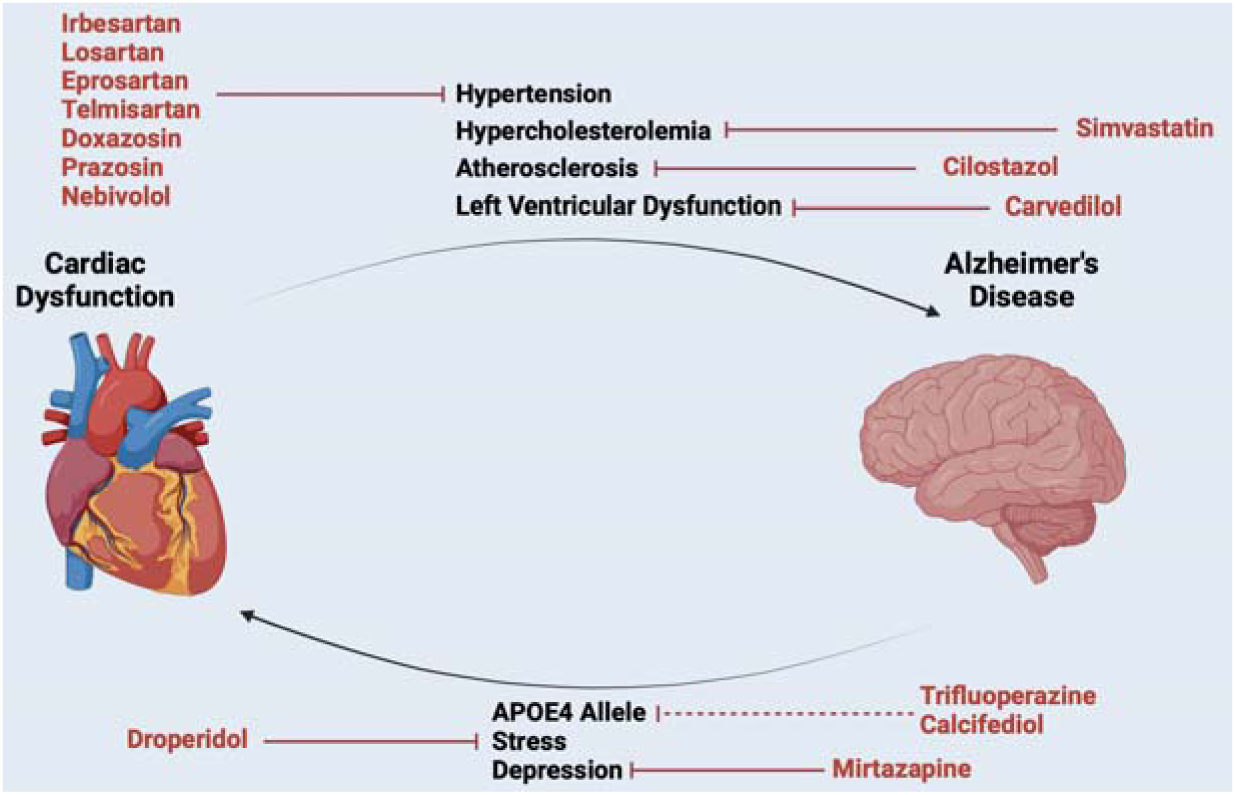
AD-CVD Axis: There is significant evidence suggesting that CVD may exacerbate neurodegeneration and that AD may cause cardiac dysfunction. Mechanistic links between hypertension, hypercholesterolemia, atherosclerosis, and left ventricular dysfunction have been observed to contribute to AD pathology, while the APOE4 allele, neural stress + inflammation, and depression are comorbid to CVD. Each heart medication explored in the study ameliorates at least one component of the shared pathology between AD and CVD. Image created in BioRender.

**FIGURE 8:**
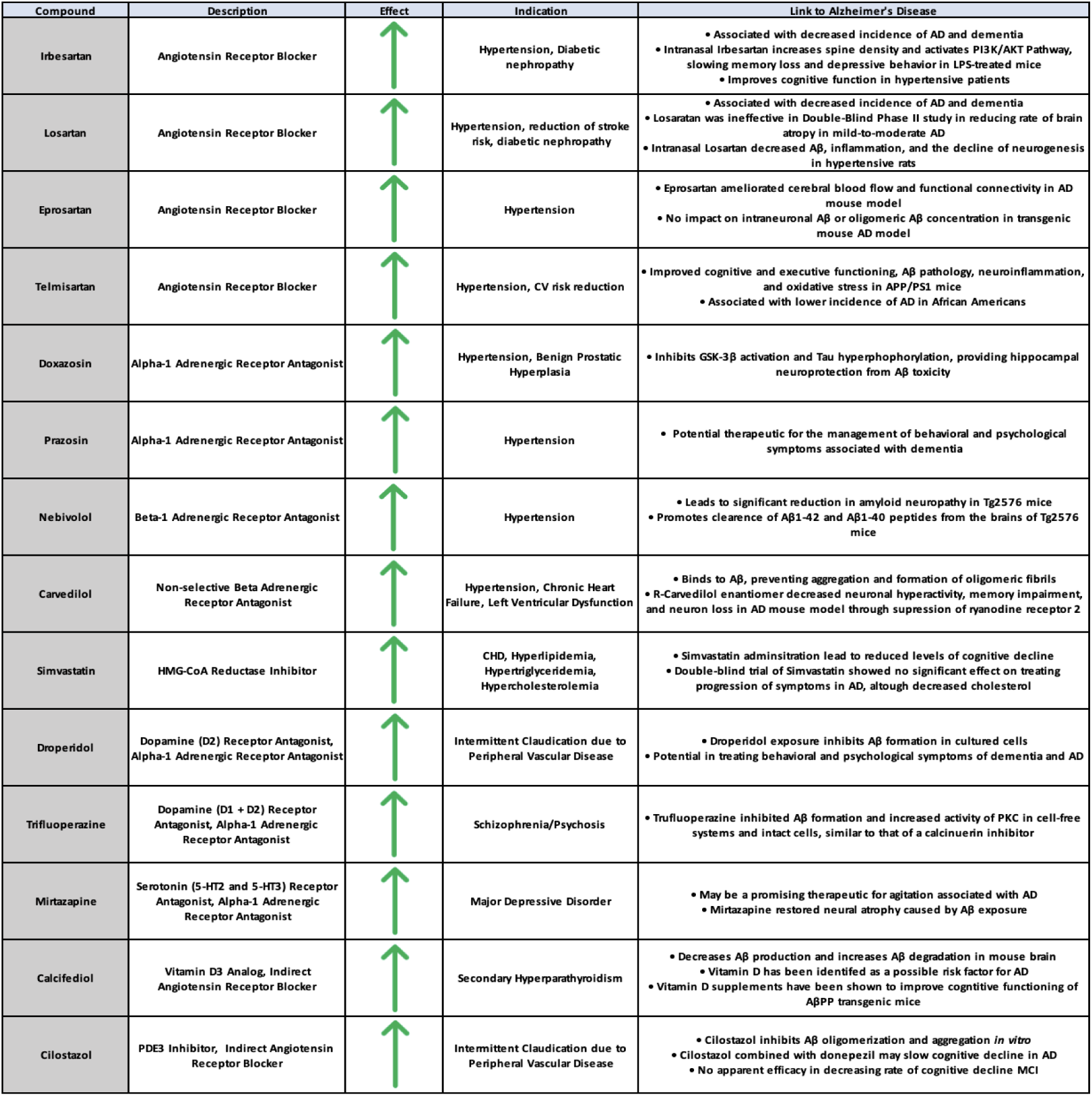
Literature Search for Heart Drugs Connection to AD: Each of the drugs explored in the study has been explored in preclinical or clinical models in treating AD and AD pathology. As denoted by an upward green arrow, all 14 of the 14 drugs may have a prospective beneficial effect in treating AD pathology based on the body of existing literature.

## DISCUSSION

### Mechanisms of action of cyclosporine and links to Alzheimer’s disease

CsA, the focus of our study, is an immunosuppressant indicated for the prophylaxis of organ rejection in kidney, liver, and heart allogeneic transplants. CsA is also approved for severe, treatment-resistant rheumatoid arthritis and plaque psoriasis. CsA forms a complex with calcineurin and cyclophilin, inhibiting the action of calcineurin {Ivery, 1999 #24}.^xi^ The CsA-calcineurin-cyclophilin complex can no longer catalyze the dephosphorylation of NFAT, which decreases T-cell activation and proliferation and regulates skeletal muscle differentiation and neurodegeneration {Mukherjee, 2011 #27}. Calcineurin signaling is primarily regulated by intracellular calcium concentrations, determined by ion pumps or release from intracellular calcium reservoirs {Park, 2019 #25}.^xii^ ^xiii^ CsA exerts its efficacy in organ rejection and skin conditions mainly through inhibiting immunocompetent T-lymphocytes and T-helper cells and blocking the production and release of lymphokines, most notably interleukin-2 {Russell, 1992 #28}.^xiv^

CsA, able to penetrate the blood-brain barrier, may have activity against neurological disorders. Longitudinal data has elucidated that calcineurin inhibitor treatment is correlated with decreased incidences of AD {Taglialatela, 2015 #5}{Silva, 2023 #29}. A recent review exploring records of calcineurin inhibitor-treated patients from a 213 million patient database (TriNetX Diamond Network) found that the general population has an increased risk of dementia compared to patients treated with calcineurin inhibitors such as tacrolimus, sirolimus, and CsA {Silva, 2023 #29}. ^xv^ The study also demonstrated that patients treated with CsA have a reduced risk of developing dementia versus patients on tacrolimus. An earlier study looking at patients in the University of Texas Medical Branch found reduced prevalence of dementia in calcineurin inhibitor-treated solid-organ transplant patients compared to the general population {Taglialatela, 2015 #5}.^xvi^ In addition, CsA has been shown to reduce biomarkers of dementia. CsA, through its ability to block the mitochondrial permeability transition pore (mPTP), halted ROS increases, mitochondrial depolarization, and restored synaptic integrity {Tapia-Monsalves, 2023 #32} affected by caspase-3 cleaved Tau (TauC3), a form of tau implicated NFT progression {Nicholls, 2017 #31}.^xvii^ ^xviii^ Other studies have demonstrated CsA’s neuroprotective properties against Aβ1-42-induced cytotoxicity {Zeng, 2013 #33}, ^xix^ and downregulation of APP expression has been observed in patients following CsA administration {Van Den Heuvel, 2004 #35}.^xx^ These observations suggest that CsA may exhibit potential as a therapeutic for AD prevention.

Aging and AD is associated with increased levels of intracellular calcium in hippocampal neurons due to oxidative stress and Aβ aggregation {Raza, 2007 #36} {Green, 2009 #37}.^xxi^ ^xxii^ We hypothesize that increased intracellular calcium leads to overactivation of calcineurin, which overactivates signaling proteins such as NFAT and GSK-3. NFAT is highly elevated in astrocytes expressing Aβ, causing neuronal death {Abdul, 2009 #177},^xxiii^ and GSK-3 is critically implicated in the progression of neuron deterioration {Rippin, 2021 #39}.^xxiv^ We believe that hyperactivity of this pathway could be a key early contributor to the pathogenesis of AD. “CsA-like drugs”, such as the ones we have identified using the Z-LaP Tracker, may have similar effects as CsA in mitigating risk of developing AD through inhibiting this pathway. However, CsA can cause serious cardiovascular, renal, endocrinological, and dermatological treatment-related adverse events. Additionally, CsA has a black box warning on its FDA label requiring experienced physicians and an equipped facility for systemic immunosuppressive therapy. Because CsA is not an ideal drug to administer chronically, we hypothesized that combining 5 µM CsA with CsA-like molecules may mitigate CsA’s side effects while still effectively targeting the pathway of interest. Moreover, the synergistic integration of CsA with another CsA-like molecule, approved to target CVD, may provide additional cardiovascular protection in an at-risk population.

### Alzheimer’s disease and cardiovascular disease

There is a well-defined axis connecting AD and CVD {Krishnamurthi, 2013 #40} {Lee, 2011 #41}{Tini, 2020 #42}.^xxv^ ^xxvi^ ^xxvii^ Factors such as hypertension, high cholesterol, obesity, and diabetes are established risk factors for CVD and major adverse cardiovascular events, and they have also been linked to an increased incidence of Alzheimer’s disease. More than 80% of AD patients show signs of CVD in autopsy {Attems, 2014 #44},^xxviii^ and AD, hypertension, and atherosclerosis are often found in tandem {Arvanitakis, 2016 #43}.^xxix^ Furthermore, ⅓ of AD-related dementia cases are due to atherosclerosis and other CVD related factors, highlighting this connection {van Gennip, 2024 #45}.^xxx^ The brain is highly vascularized {Tini, 2020 #42},^xxxi^ receiving ∼15% of total blood flow from the heart {Williams, 1989 #47},^xxxii^ and blood comprises ∼10% of the brain’s mass {Herculano-Houzel, 2009 #48} {Fantini, 2016 #50}.^xxxiii^ ^xxxiv^ The heart and brain are inextricably linked via blood perfusion, providing a clear mechanism for the transmission of pathologies. Blood perfusion is vital for the brain’s health, and ischemia causes acidosis, mitochondrial disorders, and oxidative stress in neurons {Leszek, 2021 #8}.^xxxv^ Likewise, in AD, brain hypoperfusion is a contributor to cognitive impairment and dementia {Gorelick, 2011 #52}.^xxxvi^ Furthermore, APOE4 is a significant risk factor for both AD and CVD {Stampfer, 2006 #53}, ^xxxvii^ and coronary artery disease is correlated with circulating Aβ1[40 levels {Stamatelopoulos, 2018 #54} {Stamatelopoulos, 2015 #55}{Saeed, 2023 #56}.^xxxviii^ ^xxxix^ ^xl^ Furthermore, recent studies have found common dysfunction across AD and CVD with lipid metabolism and cholesterol biosynthesis {Mathys, 2023 #57}.^xli^ In addition to cardiovascular factors exacerbating AD pathology, AD pathology can augment CVD. AD causes vascular disease, stress, and depression, all of which can advance CVD {Cortes-Canteli, 2020 #58}{Rajan, 2020 #59}.^xlii^ ^xliii^ Accordingly, there is manifold evidence of a bidirectional, self-propagating relationship between AD and CVD.

CVD often manifests up to a decade before AD {Saeed, 2023 #56},^xliv^ therefore a CVD diagnosis could mark a key time to initiate AD prevention therapy. Furthermore, drugs that treat CVD while providing neuroprotection against AD could be especially desirable. Given that we have established that CsA-like behavioral profiles identified using Z-Lap Tracker may be associated with neuroprotection, we aimed to identify drugs treating CVD that also elicit CsA-like behavior. Furthermore, we sought to identify if these drugs synergize with CsA—when combined, elicit a greater behavioral effect than when administered alone—where each drug could be administered in reduced doses, together, to achieve similar levels of efficacy. Identifying low dose, synergistic combinations would allow for lower drug exposure, reducing adverse events. This is of special importance to CsA, where reduced exposure directly increases safety and tolerability.

### Heart Drugs of Select Interest

In this study, we focused on 14 heart-targeted drugs previously identified as “CsA-like” from a high-throughput screen of the Tocriscreen FDA-Approved Drugs Library and the Cayman Chemical FDA-Approved Drug Library in 5-day-old zebrafish larvae {Gore, 2023 #6} {Del Rosario Hernandez, 2024 #7}. ^xlv^ ^xlvi^ 9 of the 14 heart drugs are FDA-approved for heart-related indications that target four common pathways. The 5 remaining drugs—droperidol, trifluoperazine, mirtazapine, calcifediol, and cilostazol— are not directly prescribed as heart medications, however, literature has suggested their high affinity towards heart-related targets. Droperidol and trifluoperazine, more commonly known as D2 antagonists, additionally block the alpha-1 adrenergic receptor {Castillo, 1995 #9} {van Nueten, 1977 #10} {Pruneau, 1984 #11} {Otani, 1989 #14}. ^xlvii^ ^xlviii^ ^xlix^ ^l^ Mirtazapine, a 5-HT2 receptor antagonist, inhibits alpha-2 adrenergic receptors {de Boer, 1995 #15}.^li^ Calcifediol (an analog of vitamin D3) serum levels are negatively correlated with the incidence of cardiovascular disease {Wang, 2016 #19} ^lii^ and calcifediol decreases ACE activity {Miller, 2022 #20}.^liii^ Lastly, cilostazol (a PDE3 inhibitor) stimulates vasodilation, inhibition of platelet activation and aggregation, and improvement of serum lipids {Weintraub, 2006 #21},^liv^ while attenuating the effects of Angiotensin II {Umebayashi, 2018 #22} {Nishioka, 2011 #23}.^lv^ ^lvi^ Consequently, all 14 drugs can be categorized as alpha-adrenergic receptor blockers, beta blockers, angiotensin receptors blockers, or statins. Our model highlights 5 of the 14 drugs, spanning all four classes, as having a potential to treat AD:

#### Statins

Simvastatin is an HMG-CoA reductase inhibitor indicated for a host of diseases related to hypercholesterolemia and hyperlipidemia. Simvastatin, a derivative of the first statin atorvastatin, inhibits the catalyzation of HMG-CoA to mevalonate, a rate-limiting step in the biosynthetic production of cholesterol. In addition to promoting nearly 50% reduction in low density lipoprotein cholesterol (LDL-C), simvastatin lowers VLDL, triglycerides, and apolipoprotein B, while increasing high density lipoprotein cholesterol (HDL-C). Long term studies have confirmed that simvastatin treatment is highly effective at improving survival in CVD patients while being “remarkably safe,” leading to few adverse events {Scandinavian Simvastatin Survival Group, 1994 #60} {Mach, 2018 #61}.^lvii^ ^lviii^

Elevated cholesterol and other lipids are a key risk factor for developing atherosclerotic CVD {Lee, 2024 #62},^lix^ a condition largely stabilized by simvastatin administration {Zhang, 2021 #63}.^lx^ It has been widely observed that there is a correlation between atherosclerosis and AD. A recent meta-analysis determined that atherosclerosis is significantly associated with AD, carotid intima–media thickness, and cognitive decline {Xie, 2020 #65}.^lxi^ Mechanistically, it has been proposed that atherosclerosis causes hypoperfusion and hypoxia in blood vessels in the brain {Zhang, 2021 #63}.^lxii^ This leads to the overproduction of Aβ, the cleavage of Aβ peptides from amyloid_β protein precursor (APP) {Garcia-Alloza, 2011 #67} {Iadecola, 2004 #68} {Koike, 2010 #69} {Li, 2009 #70},^lxiii^ ^lxiv^ ^lxv^ ^lxvi^ and a reduction in Aβ clearance, contributing to oxidative stress and neuroinflammation {Yamazaki, 2017 #71}{Chaitanya, 2012 #72}.^lxvii^ ^lxviii^ Given the shared pathophysiology, simvastatin has been explored previously as a possible therapeutic against AD.

Studies from the early 2000s have shown simvastatin’s significant ability to reduce intracellular and extracellular levels of Aβ42 and Aβ40 in hippocampal neurons and mixed cortical neurons {Fassbender, 2001 #73},^lxix^ and more recent studies have shown simvastatin reduce sAβ42 in yeast, neuroblastoma cell lines {Ostrowski, 2007 #74},^lxx^ and in human brain deposits {Nabizadeh, 2023 #75}.^lxxi^ Furthermore, longitudinal and epidemiological studies have substantiated that simvastatin administration is correlated to a strong reduction in the incidence of AD and an increase in cognitive function {Reiss, 2005 #76}{Petek, 2023 #77}.^lxxii^ ^lxxiii^ These lines of evidence suggest that simvastatin may be an effective treatment for AD. However, a 2011 Phase II trial exploring simvastatin vs. placebo in subjects with probable AD showed that simvastatin was unable to slow cognitive decline according to the Alzheimer’s Diseases Assessment Scale-cognitive portion (ADAS-Cog) {Sano, 2011 #78}.^lxxiv^ These results are consistent with a randomized controlled trial of atorvastatin in mild to moderate AD, where the treatment was ineffective in improving cognition on the same scale {Feldman, 2010 #79}.^lxxv^ However, the participants in the simvastatin trial had an average age of approximately 75 years, which we believe contributed to the drug’s observed lack of efficacy. In contrast, longitudinal studies showing a decrease in AD incidence with simvastatin involved younger populations, with a median age of 65, who had been taking the drug for at least three years {Torrandell-Haro, 2020 #80}.^lxxvi^ This indicates that earlier administration of simvastatin in the progression of AD might enhance its effectiveness.

Meanwhile, simvastatin also demonstrates significant action in the calcineurin-NFAT signaling pathway. A recent publication revealed that simvastatin is a potent inhibitor of Hsp90 (heat shock protein 90), an abundant molecular chaperone involved in many cell signaling pathways {Jackson, 2013 #81}.^lxxvii^ Hsp90 is required for activating calcineurin and c-Raf, and therefore simvastatin administration was found to decrease calcineurin and c-Raf expression {Jackson, 2013 #81}.^lxxviii^ This provides a mechanistic connection between the action of simvastatin and CsA.

Our study indicates high levels of synergy between simvastatin and CsA. Simvastatin and its 5 µM combination with CsA demonstrated a CsA-like behavioral profile in our Z-LaP Tracker model. This is classified by increases in baseline activity and activity when exposed to visual stimuli, large decreases in habituation, increased excitability, and a decreased optomotor response. Our lab has previously associated this behavioral profile with compounds that could potentially be protective against neurodegeneration and AD {Gore, 2023 #6}{Del Rosario Hernandez, 2024 #7}.^lxxix^ ^lxxx^ Our IPA analysis showed the highest levels of predicted inhibition of AD with simvastatin treatment, especially when compared to other drugs in our model explored. 5 µM simvastatin + 5 µM CsA elicited the most CsA-like behavioral profile (even when compared to 5 µM CsA alone), with an R value of 0.86. Furthermore, 5 µM simvastatin + 5 µM CsA led to a 22% increase in the magnitude of behavior in Z-LaP Tracker model compared to a higher dose of simvastatin alone, indicating that simvastatin and CsA potentiate one another. This is consistent with human pharmacokinetic data, as the American Heart Association in 2016 issued a statement in 2016 that the combination of simvastatin and CsA can lead to 6-8x increases in the AUC of simvastatin {Wiggins, 2016 #84}.^lxxxi^ While this could be potentially harmful, combining lower doses of each drug has yet to be explored. According to our cluster analysis, simvastatin and CsA elicited the most similar behavioral profiles. This evidence suggests that combining low doses of simvastatin and CsA could double as an effective CVD medication *and* a preventative treatment for AD.

#### Angiotensin Receptor Blockers

Irbesartan is a non-peptide angiotensin II competitive antagonist indicated for hypertension and diabetic nephropathy in hypertensive patients with type 2 diabetes, elevated serum creatinine, and proteinuria {Darwish, 2021 #87}.^lxxxii^ Through its action at the Angiotensin AT1 Receptor, irbesartan normalizes vasoconstriction and aldosterone-secretion stimulated by angiotensin II. Irbesartan has >8500-fold greater affinity for AT1 receptors than AT2 receptors and has been found to have no appreciable effect on ACE, renin, or any other cardiovascular related receptor, ion or hormone.

Given hypertension’s link to AD, irbesartan has been implicated as a potential therapeutic for Alzheimer’s disease. A recent study found that intranasal irbesartan increased dendritic spine density, decreased synaptic dysfunction, and activated the P13K/AKT pathway, leading to a decrease in memory loss in LPS-treated mice {Gouveia, 2024 #88}.^lxxxiii^ This suggests that irbesartan could be repurposed as a preventative treatment against AD. To this extent, a review of US VA data concluded that patients on irbesartan have significantly lower incidences of dementia than the general population (HR 0.84, P < 0.001), and this relationship was observed in a dose-dependent manner {Li, 2010 #89}.^lxxxiv^ Behaviorally, hypertensive patients treated with irbesartan experienced positive effects on long-term memory and psychomotor vitality compared to baseline {Hao, 2024 #90}.^lxxxv^

Meanwhile, literature has suggested that irbesartan significantly reduces calcineurin expression and calcium-calcineurin signaling {Jiang, 2006 #91} {Shang, 2011 #92}.^lxxxvi^ ^lxxxvii^ Although through a different mechanism, irbesartan like CsA, inhibits calcineurin. We suggest that irbesartan’s anti-neurodegenerative profile may be bolstered by its ability to decrease calcineurin. Irbesartan, and 5 µM irbesartan + 5 µM CsA, formed a significant subcluster (Subcluster B, Figure 5) with CsA and demonstrated highly CsA-like behavior in our Z-LaP Tracker model. This was typified by highly pronounced increases in 1-hour activity and activity during visual stimuli. Furthermore, 5 µM irbesartan + 5 µM CsA synergized to elicit a 53% larger response in magnitude than irbesartan alone. Of note, 5 µM CsA + 5 µM irbesartan caused an additional 3 changes in behaviors to be significantly different compared to irbesartan alone. In addition, our correlation matrix revealed that 5 µM irbesartan + 5 µM CsA and irbesartan alone have highly correlated behavioral values. This corroborates that CsA and irbesartan may target similar pathways and have synergistic effects. We highlight irbesartan as a drug of interest for future studies.

Cilostazol is a quinolinone derivative that inhibits phosphodiesterase III (PDE3). PDE3 inhibition leads to the suppression of cAMP degradation, increasing circulating cAMP and inhibiting platelet aggregation and vasodilation. Accordingly, cilostazol is FDA approved for the treatment of intermittent claudication, a condition where lack of oxygen delivered to the leg muscles causes pain, cramping, and discomfort. Cilostazol, as with other PDE3 inhibitors, contains a black box warning, contraindicated in patients with a history of heart failure. Given cilostazol’s anti-hypertensive properties, it has been explored as a potential inhibitor of angiotensin. Multiple studies have confimred that cilostazol administration suppresses angiotensin II activity, decreasing endothelial cell apoptosis and dysfunction {Shi, 2016 #119} ^lxxxviii^ and angiotensin II-induced aortic aneurysm {Umebayashi, 2018 #22}.^lxxxix^ Consequently, although cilostazol does not directly target angiotensin, we are classifying cilostazol with the other angiotensin receptor blockers for our data analysis.

There is a significant body of evidence suggesting cilostazol may have efficacy as a treatment against AD. Preclinical models have demonstrated that cilostazol administration can sequester Aβ-induced neurotoxicity, stimulate clearance of soluble Aβ, and reverse cognitive impairment {Ono, 2019 #121}.^xc^ Furthermore, a case-control human study revealed that cilostazol may be protective against cognitive decline in AD {Tai, 2017 #122}.^xci^ Similar to the other selected drugs, cilostazol also has relevance in the calcineurin-NFAT pathway. Cilostazol mitigates production of various inflammatory molecules transcribed by activation of this pathway, including nitric oxide, prostaglandin E2, proinflammatory cytokines, interleukin-1 (IL-1), and tumor necrosis factor-α {Jung, 2010 #123}.^xcii^ Accordingly, in our study, 5 µM cilostazol + 5 µM CsA led to the greatest number of new significantly different behaviors compared to cilostazol alone, driven by significant increases in P15, scoot, and burst behaviors. This indicates extraordinarily high synergy between cilostazol and CsA. Interestingly, two past studies have evaluated adjunctive treatment of CsA and cilostazol. Rat models have revealed that cilostazol reduced CsA-associated renal ischemia-reperfusion injury {Gokce, 2012 #124},^xciii^ and reduces neointimal hyperplasia following vascular injury {Badiwala, 2010 #125}.^xciv^ This dual benefit suggests a perfect synergistic combination, potentially offering improved efficacy outcomes with reduced side effects. Our results and review highlight cilostazol as a drug of interest for AD treatment, especially in combination with CsA.

#### Alpha Adrenergic Antagonists

Doxazosin, a quinolone derivative, is a competitive alpha1-antagonist targeting the postsynaptic alpha1 receptor. Doxazosin is FDA approved for two indications: hypertension and the treatment of Benign Prostatic Hyperplasia (BPH). Doxazosin has a long half-life, about 16-30 hours, and therefore only has to be dosed once daily {Smith, 2016 #126}.^xcv^ Alpha1 receptors are predominantly found in vascular smooth muscle where they regulate arteriolar resistance and venous capacitance {Reid, 1986 #127}.^xcvi^ Doxazosin administration prevents the binding of norepinephrine to the alpha1-adrenergic receptor, relaxing smooth muscle cells, reducing peripheral vascular resistance and lowering blood pressure {Remaley, 2007 #128}.^xcvii^ Studies have also shown that doxazosin has an appreciable effect blocking other signaling pathways such as intracellular calcium influx via voltage-dependent and - independent calcium channels and the activation of phospholipase A2 {Insel, 1989 #129}.^xcviii^ Consequently, in addition to blocking the alpha1-adrenergic receptor, doxazosin leads to substantial decreases in intracellular calcium concentrations, suppressing calcineurin signaling, similar to CsA.

Research has suggested that doxazosin may have potential as an AD therapeutic. *In vitro*, doxazosin has demonstrated a robust ability to reduce tau hyperphosphorylation and block GSK-3ß activation, mitigating Aß toxicity {Coelho, 2019 #130}.^xcix^ Subsequent studies have presented that doxazosin significantly increases BDNF and Akt kinase activity in the cerebral cortex, correlating to neuroprotection {Mohamed, 2024 #131}.^c^ Furthermore, given the relationship between hypertension and AD, there could be other mechanistic pathways by which doxazosin contributes to resilience against AD pathology. There have yet to be any human trials exploring the efficacy of doxazosin in treating AD.

Our Z-LaP Tracker model revealed high levels of synergy between doxazosin and CsA. While doxazosin alone leads to significant increases in excitability and 1-hour activity (both “CsA-like” behaviors), 5 µM doxazosin + 5 µM CsA added 3 additional significant behaviors and a 20% total increase in behavioral magnitude. Importantly, the combination led to a significant *decrease* in habituation behavior, a key “CsA-like” characteristic. In our cluster analysis, doxazosin and 5 µM doxazosin elicited the most CsA-like behavioral profile: not only did 10 µM doxazosin and 5 µM doxazosin cluster within Subcluster A, but they also clustered directly in between 5 µM CsA and 10µM CsA. Interestingly, in our correlation matrix, doxazosin and its 5 µM combination had positive correlation with 5 µM CsA but a lesser relationship with 10µM CsA. This is likely due to differences in edge behavior. Furthermore, IPA’s Path Explorer model revealed a high level of overlap between doxazosin and AD, where upregulation of doxazosin leads to high predicted inhibition of AD pathology. We highlight doxazosin as a molecule of special interest as a combination with CsA or other CsA-like drugs for the treatment of AD.

#### Beta Adrenergic Antagonists

Nebivolol is a beta-adrenergic blocker used to lower blood pressure in hypertension. Nebivolol is cardioselective, only blocking beta1-adrenergic receptors located in cardiac tissue. According to its FDA label, there are five mechanisms by which nebivolol treats hypertension: through decreasing heart rate, decreasing myocardial contractility, decreasing outflow, suppressing renin activity, and vasodilation. Nebivolol does not have any affinity to alpha1-adrenergic receptors at clinically relevant doses but is unique amongst other beta blockers due to its ability to stimulate nitric oxide-induced vasodilation {Weiss, 2006 #160}.^ci^

Nebivolol increases nitric oxide (NO) synthase through beta-3 agonism from the endothelium, inducing vasodilation {Ågesen, 2019 #161}.^cii^ NO is protective against reactive oxygen species-mediated oxidative stress and target organ damage, therefore nebivolol’s efficacy in hypertension may extend to other disorders impacted by oxidative dysfunction such as AD {Coats, 2017 #162}.^ciii^ While nebivolol and CsA do not target the same pathway, they both decrease the transcription of inflammatory factors {Barroso, 2022 #163} and therefore have mechanistic similarities.^civ^ In a rat model of cerebral ischemia/reperfusion injury, nebivolol is able to alleviate oxidative stress through its regulation of eNOS and iNOS {Heeba, 2012 #164}.^cv^ Interestingly, nebivolol can penetrate the BBB and appreciably reduce Aβ neuropathy in the brain and plasma Aβ levels in Tg278 mice {Wang, 2013 #165}.^cvi^

Our results for nebivolol are especially interesting. Although nebivolol and CsA have different mechanisms of action, they elicit highly similar behavioral profiles in our Z-LaP Tracker model: the 5 µM nebivolol + 5 µM CsA combination was grouped in Subcluster B. Furthermore, despite the negative values in Figures 2A and 2B, 5 µM nebivolol + 5 µM CsA led to substantial increases in the magnitude of activity and vision behaviors. Although Figures 2A and 2B show decreases in the number of significant behaviors and behavioral magnitude, we observed high levels of synergy between nebivolol and CsA in several key behaviors. This indicates to us that nebivolol and CsA may be highly complementary to one another. The correlation matrix revealed high levels of correlation between CsA and nebivolol and CsA and the 5 µM nebivolol combination. IPA’s pathway analysis feature also predicts moderate levels of inhibition of AD pathology with nebivolol upregulation. Given the body of evidence, we highlight nebivolol and the 5 µM nebivolol combination therapies of key interest in protecting against neurodegeneration.

## CONCLUSION

Our study’s aim was to identify FDA approved compounds to treat CVD that elicit in our model a “CsA-like” behavioral profile, indicating potential neuroprotection against AD and dementia. We explored which of these 14 CVD drugs, when combined with CsA, synergized to show more significant behavioral changes compared to when administered alone. Our model, Z-LaP Tracker, highlighted 5 of the 14 drugs to be most “CsA-like” *and* to positively synergize with CsA: simvastatin, irbesartan, cilostazol, doxazosin, and nebivolol. These combinations may have promise to protect against AD while also treating CVD. Drugs across all four classes explored (alpha antagonists, beta blockers, angiotensin receptor blockers, and statins) demonstrated viability and synergy across our experiments. This suggests that treatments across various manifestations of CVD could be leveraged to confer neuroprotection. In the future, our lab seeks to utilize Western blot and RNA-sequencing to elucidate the molecular mechanisms underpinning the behaviors elicited by these drugs and to confirm their action in the calcineurin pathway.

## METHODS

### Animal Handling & Husbandry

The research outlined in this study has been conducted in accordance with federal regulations and guidelines for the ethical and humane use of animals and has been reviewed and approved by Brown University’s Institutional Animal Care and use committee (IACUC). All experiments in this study were performed on 5 days post fertilization (dpf) zebrafish larvae obtained from genetically diverse outbred strain of adult wild-type zebrafish (*Danio rerio*). The adult zebrafish are maintained in a 14-h light, 10-h dark cycle environment in a Marineland Vertical Aquatic Holding System containing 15- and 30-gallon tanks. Fish are bred and fed with Gemma Micro 300 and frozen brine shrimp daily. At 0-5 dpf, the zebrafish embryos/larvae are maintained in 28.5 °C 2L culture trays in egg water (60 mg/L sea salt (Instant Ocean) and 0.25 mg/L methylene blue in deionized water (pH 7.2)), on a 12-h light/12-h dark cycle. By 5 dpf, larvae are able to display various complex behaviors and do not require external food source, consuming nutrients from the yolk sac {Clift, 2014 #179}. After the completion of the behavioral explements, tested larvae were euthanized by rapid chilling followed by immersion in a bleach solution (1 part bleach, 5 parts tank water) for 30 minutes each.

### Pharmacological Treatment

Zebrafish larvae five days post fertilization (dpf) were exposed to 14 small-molecules identified from the Cayman Chemical FDA-approved Drug Library and the Tocriscreen Small-Molecule Library. These 14 compounds represent the only cardiovascular targeted drugs identified as “CsA-like” from our previous studies. Each compound was diluted to 10 mM stocks in dimethyl sulfoxide (DMSO), and later diluted with egg water to 10 µM when administered alone and 5 µM when in combination with CsA. Zebrafish larvae are individually placed in an opaque well in a 96-well ProxiPlate (PerkinElmer 6006290) containing 100 µL of the treatment or control solution, where they are exposed starting 3 hours prior to the behavioral assay. The larvae remained in the same treatment solution during imaging. The control groups were larvae in egg water and in 1 µL/mL DMSO.

### Behavioral Assay & Z-LaP Tracker

We employed Z-LaP Tracker to analyze how drug exposure affected zebrafish behavior. The behavior of zebrafish larvae was examined using a custom-built imaging system as described previously {Gore, 2023 #6}. Up to 4 plates can be imaged at once, each holding up to 96 larvae in a sound, light, and temperature-controlled environment. Images of the larvae were captured every 6 seconds using a high-resolution camera. Visual and acoustic stimuli were presented to the larvae as a 3-hour Microsoft PowerPoint presentation, as described previously {Gore, 2023 #6}. The PowerPoint includes slow- and fast-moving colored lines, and acoustic stimuli of different intervals.

As previously described {Gore, 2023 #6}, Z-LaP Tracker can detect 25 different behavioral parameters through automated tracking of a larva’s eyes and yolk: 1-h activity = % of time that the larvae move during the first hour of imaging; P15 = % of time that the larvae move during period 15 (140-150 min); Hab = Change in activity in response to 1s sound intervals, period 17; S = change in activity in response to 20s sound intervals; E = change in activity in 1s acoustic stimuli, period 16-17; R = optomotor response to red moving lines; G = optomotor response to green moving lines; B = optomotor response to blue moving lines; FR = optomotor response to faster moving red lines; RGB = combined optomotor response; Sc-1h = % of time in first hour moving in slow/short swimming pattern (scoot); Sc-V = % in scoot behavior during moving lines; Bu-1h = % of time in first hour moving in quick/long swimming pattern (burst); Bu-V = % in burst behavior during moving lines; Ed-1h = % of time located in edge of well in first hour; Ed-V = % of time located in edge of well during moving lines; Cw-1h = % of time orientation is clockwise during first hour; Cw-V = % of time orientation is clockwise during moving lines; Or = % of time with upward orientation; Turn-1h = change in larvae’s position angle during first hour; Tabs-1h = absolute change in position angle during first hour. The behaviors are summarized in an an excel template, which combines and summarizes the results of multiple experiments {Gore, 2023 #6}.

### Statistical Analysis

Statistical tests and data graphs were generated using Microsoft Excel 2016 and R. We used the non-parametric Welch’s unequal variance t-test with subsequent Bonferroni correction for multiple comparisons due to the nature of our data. We compared 30 individual and combination treatments to the DMSO vehicle controls and differences were considered significant when p < 1.67 × 10^-3^ (0.05/30), p < 3.33 × 10^-4^ (0.01/30), or p < 3.33 × 10^-5^ (0.001/30). Correlation analysis was performed using the corrplot package in R.

### Cluster Analyses

Behavioral profiles were generated from changes in larval activity, habituation, startle response, excitability, and optomotor response compared to DMSO vehicle controls. Further analyses of these profiles was performed in Microsoft Excel using previously described templates {Gore, 2023 #6}. Cluster 3.0 and TreeView were used for hierarchical cluster analysis and visualization, respectively. Hierarchical clustering was performed using Euclidean distance as a similarity metric, and complete linkage.

### IPA Analysis

QIAGEN’s Ingenuity Pathway Analysis (IPA) was used to explore direct and indirect links between the effects of the selected heart drugs and AD. We created four separate “My Pathways” based on the category of the drug: angiotensin receptor blocker, beta blocker, alpha adrenergic receptor antagonist, and HMG CoA Reductase Inhibitor. Using the “Path Explorer” feature, we generated direct and indirect relationships between each molecule and AD using the “Ingenuity Knowledge Base”. Additionally, we generated an overlay indicating activation or inhibition of pathway components and subsequent connections using the “Molecule Activity Predictor (MAP)” tool.

## Acknowledgment

This work was supported by the NIH, NIGMS R01GM136906 (RC)

## Supplementary

**FIGURE 9:**
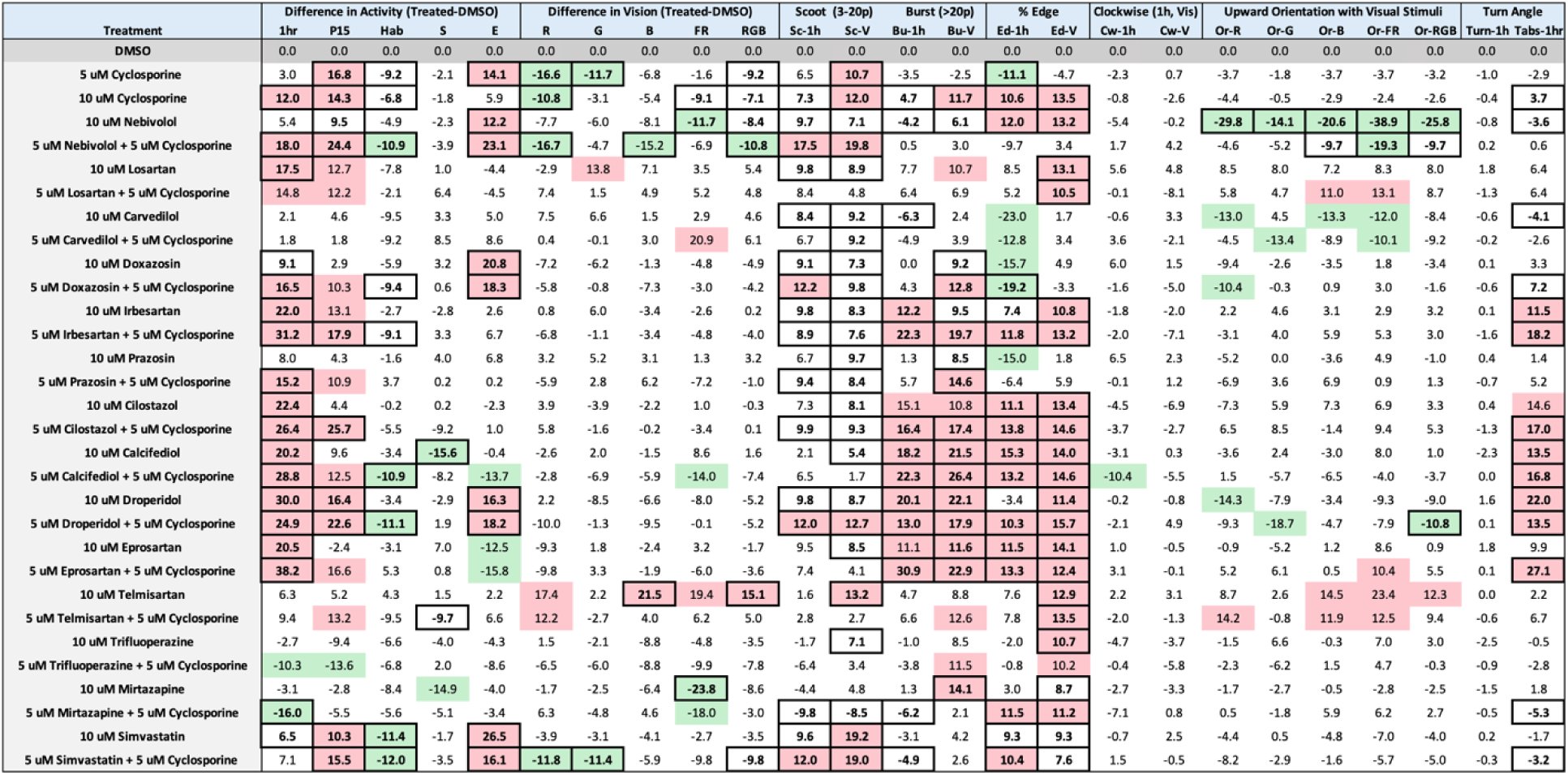
25 Behavioral Measures Calculated using Z-LaP Tracker: The behavioral changes associated with 30 experimental treatments when compared to a DMSO control. Each value represents the percentage point difference between the active treatment and DMSO control. A 10%-point increase in a behavioral measure is illustrated by red boxes, and a 10%-point decrease by green boxes. Significant changes in a behavior parameter are denoted by a bolded and boxed cell (p < 1.67 × 10^-3^, correction for multiple comparisons [0.05/30]). Over 2300 larvae were tested, and the smallest n per arm was 30.

